# Consequences of breed formation on patterns of genomic diversity and differentiation: the case of highly diverse peripheral Iberian cattle

**DOI:** 10.1101/466821

**Authors:** Rute R. da Fonseca, Irene Ureña, Sandra Afonso, Ana Elisabete Pires, Emil Jørsboe, Lounes Chikhi, Catarina Ginja

## Abstract

**Background:** Iberian primitive breeds exhibit a remarkable phenotypic diversity over a very limited geographical space. While genomic data are accumulating for most commercial cattle, it is still lacking for these primitive breeds. Whole genome data is key to understand the consequences of historic breed formation and the putative role of earlier admixture events in the observed diversity patterns.

**Results:** We sequenced 48 genomes belonging to eight Iberian native breeds and found that the individual breeds are genetically very distinct with F_ST_ values ranging from 4 to 16% and have levels of nucleotide diversity similar or larger than those of their European counterparts, namely Jersey and Holstein. All eight breeds display significant gene flow or admixture from African taurine cattle and include mtDNA and Y‐chromosome haplotypes from multiple origins. Furthermore, we detected a very low differentiation of chromosome X relative to autosomes within all analyzed taurine breeds, potentially reflecting male‐biased gene flow.

**Conclusions:** Our results show that an overall complex history of admixture resulted in unexpectedly high levels of genomic diversity for breeds with seemingly limited geographic ranges that are distantly located from the main domestication center for taurine cattle in the Near East. This is likely to result from a combination of trading traditions and breeding practices in Mediterranean countries. We also found that the levels of differentiation of autosomes *vs* sex chromosomes across all studied taurine and indicine breeds are likely to have been affected by widespread breeding practices associated with male-biased gene flow.

## Background

The biological resources of the Mediterranean sub-region of the Palaearctic include a diversity of domesticated animals [1] comprising 53 officially recognized local breeds of taurine cattle (*Bos taurus*) in the Iberian Peninsula alone (Table S1). Taurine cattle are thought to have been domesticated by Neolithic farmers from *B. primigenius* populations in the Fertile Crescent around 10,000 years [2], and have since diversified into more than 1,000 breeds [3]. Cattle genomes have been shaped not only by human-driven selection, but also by genetic bottlenecks associated with migrations from the origin of domestication, adaptation to different agro-ecological areas and a more strict division of animal populations into breeds led by Europeans since the 18^th^ century [3]. Furthermore, multiple events of introgression have been proposed to have influenced European cattle breeds: i) ancestral hybridization with European populations of *B. primigenius* [4–9] (extinct in Europe since the 17^th^ century [9]); ii) introgression from African taurine cattle [10]; iii) introgression from non-taurine sources such as indicine breeds (*Bos indicus*, the humped cattle type resulting from a separate domestication event in the Indus valley [11]) [10,12]. Such wide-spread gene-flow resulted in complex patterns of admixture and the difficulty in sometimes establishing whether a breed represents the taurine populations that were originally associated with a specific geographic region [10] and could explain the high levels of genetic diversity relative to other domesticated species [12].

Currently, there are two broad groups of cattle breeds, those under intensive management with strong specialization in dairy or meat phenotypes (such as the commercial transboundary Holstein, Charolais, Limousine, and more recently Angus), and the so-called “primitive” breeds, traditional cattle with a low dependence from external inputs that make use of naturally available food resources. Iberian native cattle are found in diverse agro-ecological systems including coastal, mountain, and lowland arid environments (Fig. 1A). Inheritable traits of these cattle have been modified at different times by the various cultures that inhabited this territory, and breeds are often defined based on morphological traits such as coat color, as well as horn size and body shape.

**Figure 1.**
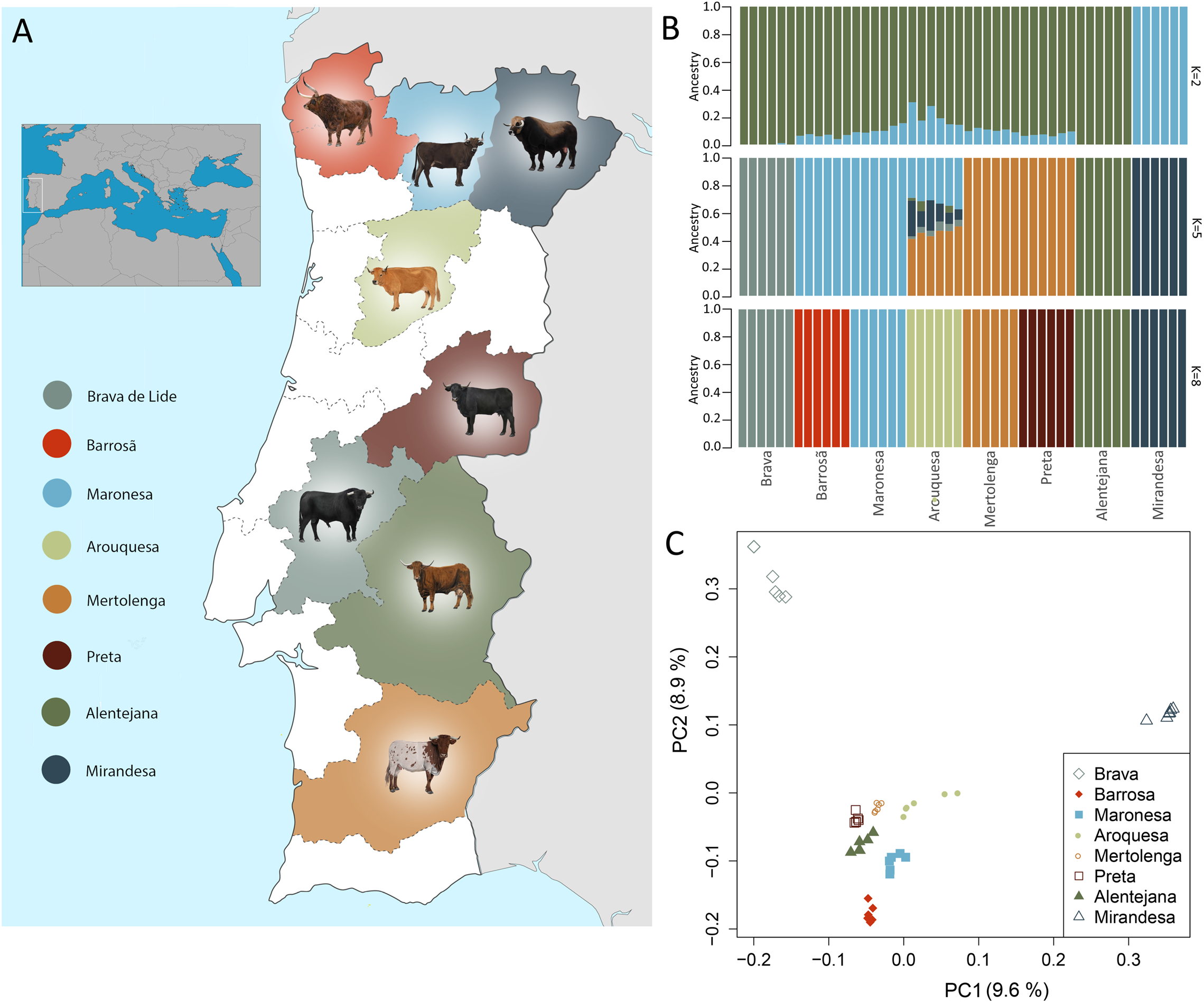
**A)** Geographical distribution of the eight Iberian native breeds. **B)** Population structure plot determined by NGSadmix shows consistency with breed denomination; each individual is represented by a stacked column of the 2, 5 and 8 proportions (other K values in Figure S3). **C)** Reproductive isolation of the Mirandesa and Brava breeds relative to the others is clear in the principal component analysis done with PCAngsd; variance explained by each component is shown in parenthesis (other components are in Figure S2).

Recently, the Food and Agriculture Organization (FAO) has warned that about 67% of the Iberian cattle breeds are at risk as many of these have less than 1,000 breeding females and/or less than 20 breeding males [3], which reinforces the need for a continued conservation strategy. The complex origin of the Iberian primitive breeds is reflected in their high diversity in Y-chromosome haplotypes, including the major taurine Y1 and Y2 haplogroups [13,14] and unique patrilines [15], as well as distinct maternal lineages, i.e. common European T3-matrilines along with more distinct Q-haplotypes [14,16,17], and a strong influence of T1-lineages of African origin [18]. This higher diversity relative to their European counterparts is quite notable, given the geographic distance of this territory from the presumed Near-Eastern center of domestication [4,13,14,19,20]. This renders Iberian cattle a great example for investigating the genomic impact of the intricate processes of cattle diversification both regarding the last 200 years of specific breed formation and the putative earlier admixture events.

To uncover genome-wide patterns of diversity associated with the formation of primitive cattle breeds, we sequenced the genomes of 48 individuals belonging to eight breeds of native Iberian cattle (Fig. 1A). Their breed denominations have been shown to agree with population structure inferred from microsatellites [16,19–21]. Noteworthy, no clear structure is recovered when using genotypes determined with the Illumina Bovine High-Density 777k SNP BeadChip in the context of European cattle [4], likely a result of ascertainment bias as Iberian breeds were not included in the discovery panel of the genotyping assay. This reinforces the need for full genome data to accurately determine genetic diversity and measure population differentiation [22].

We confirm that there is a clear genetic distinction between Iberian cattle breeds. In addition, we demonstrate that breed management and associated demographic processes had profound effects on genomic diversity and resulted in unusual patterns of genetic differentiation for autosomes *vs* sex chromosomes. We further describe genome-wide diversity and introgression in Iberian breeds in relation to 60 previously published taurine (*B. taurus*) and zebu (*B. indicus*) cattle genomes from Europe, Africa and Asia, and sequence data from one European aurochs (*B. primigenius*). We confirm that gene flow has occurred between African taurine and Iberian breeds. Overall, we show how whole-genome data are important for uncovering specific patterns related to recent events in breed formation and management, and provide the ground for future studies on the singularity of locally adapted European cattle breeds.

## Results and Discussion

The 48 Iberian cattle genomes and the previously published shotgun resequencing data from 60 additional individuals including taurine and indicine cattle (Additional file 1: Table S3) were mapped with BWA mem to three reference genomes: genome version UMD_3.1.1 (bosTau8) [23], genome version Btau_4.6.1 (bosTau7; contains an assembled Y-chromosome) [23] and to the outgroup wild yak (*B. mutus*) [24]. Details on the quality-based read trimming and filtering steps are included in the Methods section. Sequencing error rates for all 48 samples are below 0.2% (Additional file 1: Figure S1).

### Signatures of breeding in the population structure and genetic differentiation of Iberian cattle breeds

Population structure and individual ancestry were investigated with NGSadmix, which does not require definition of the exact genotypes thus is adequate for low-depth sequencing data [25]. Setting the number of expected clusters to eight (the number of breeds) resulted in the assignment of each individual to the source breed (Fig. 1B) while assuring convergence of the method. This level of genetic homogeneity within Iberian cattle populations is also observed in the results of the principal components analyses (Fig. 1C). The first two PCs explain 10% and 9% of the total variation and show the high differentiation of Mirandesa and Brava. Mirandesa in fact appears as an independent cluster when the number of ancestral *K* populations is set to two (Fig. 1B), and Brava individuals become a separate cluster when *K* = 4 (Additional file 1: Figure S2). Both observations are expected to result from genetic drift due to drastic demographic changes: in the 1970s, Mirandesa was raised in a vast area of the Portuguese territory with over 200,000 animals [26] and since has suffered a significant reduction in population size with less than 6,000 breeding females registered in the herdbook in 2017 (http://www.fao.org/dad-is/browse-by-country-and-species/en/); Brava has historically been reproductively isolated from other breeds living in semi-feral conditions for the main purpose of its use in bullfights [26]. PCs 3 and 4 separate Alentejana and Preta from the remaining Portuguese native breeds, whereas Maronesa, Barrosã and Mertolenga are separated by PCs 5 and 6 (Additional file 1: Figure S3).

Recent crossbreeding involving Arouquesa cattle is revealed in it being the last to form a discrete cluster, showing contributions from the other populations until *K* = 7 (Fig. 1B and Additional file 1: Figure S2). This is consistent with an analysis of microsatellite loci, which showed Arouquesa as having the lowest mean genotype membership proportions [19]. This breed is mostly raised in a region located south of the Douro river in the district of Viseu (Fig. 1A), bordering the area of production of Maronesa and in remote times also of the once abundant Mirandesa cattle. Arouquesa has also historically been crossbred with the latter to produce the highly valued “*vitela de Lafões*”, a meat product certified by the European Union with Protected Geographical Indication, and so admixture is intrinsically linked to its history. Another breed showing high heterogeneity was Mertolenga (Additional file 1: Figure S2), one of the most phenotypically diverse Iberian native breeds, with its three distinct coat color phenotypes mostly raised in separate herds [19].

We assessed the levels of genetic differentiation between breeds by calculating the fixation index (F_ST_).In general, we observed high levels of differentiation (average 9%), even when admixture has occurred, which precludes the use of Iberian cattle as a single evolutionary unit. Consistent with their higher heterogeneity, the breed pair Arouquesa/Mertolenga had a low F_ST_ value of 0.06. The highest F_ST_ corresponded to the pairwise comparison of Mirandesa and Alentejana (F_ST_ = 0.16) and the lowest F_ST_ values were obtained for Preta *vs* Mertolenga (F_ST_ = 0.04) (Table 1).

**Table 1.** F_ST_ values between the eight Iberian breeds. The highest value is shown in bold and the lowest in italic.

|  | Alentejana | Arouquesa | Barrosã | Brava | Mertolenga | Mirandesa | Maronesa |
| --- | --- | --- | --- | --- | --- | --- | --- |
| **Arouquesa** | 0.10 |  |  |  |  |  |  |
| **Barrosã** | 0.12 | 0.06 |  |  |  |  |  |
| **Brava** | 0.13 | 0.08 | 0.09 |  |  |  |  |
| **Mertolenga** | 0.09 | 0.06 | 0.06 | 0.07 |  |  |  |
| **Mirandesa** | **0.16** | 0.08 | 0.11 | 0.13 | 0.11 |  |  |
| **Maronesa** | 0.12 | 0.06 | 0.06 | 0.09 | 0.06 | 0.11 |  |
| **Preta** | 0.08 | 0.05 | 0.06 | 0.05 | *0.04* | 0.09 | 0.06 |

### Iberian genetic variation in the context of taurine and zebu cattle diversity

When compared to publicly available genomes [27] (database information in the Materials section) of taurine (European Holstein, Angus and Jersey, and African N’Dama) and African indicine cattle (Ogaden, Kenana and Borana), Iberian breeds are clearly assigned by NGSadmix [25] to the European cluster (Fig. 2A) with a slight suggestion of African taurine admixture at *K* = 3 for autosomal data. As observed previously [27], at *K* = 3 the clusters observed represent European taurine, African taurine and African indicine ancestries.

**Figure 2.**
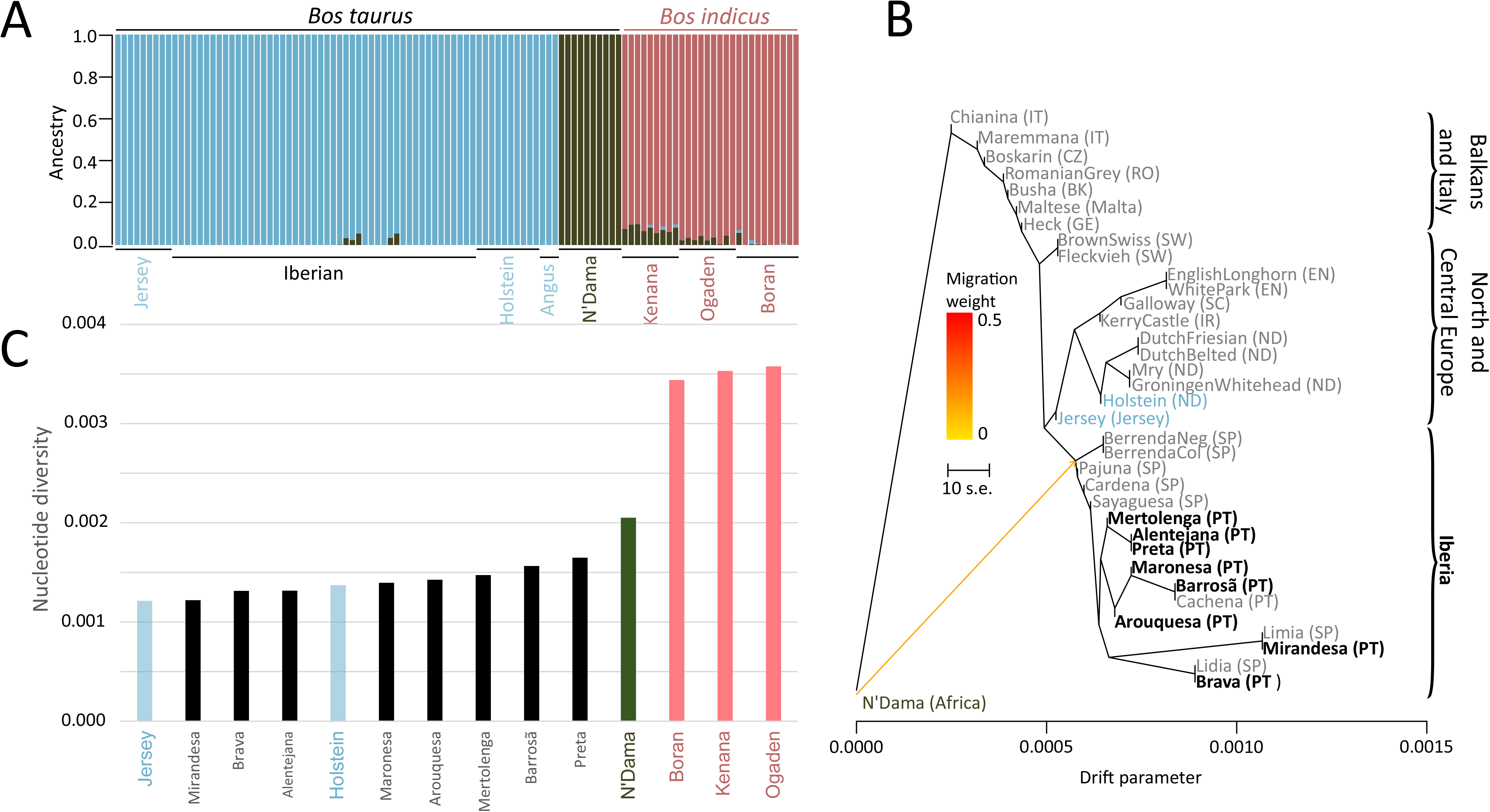
**A)** Population structure using 108 individuals at K=3 clearly divides the European taurine (blue), African taurine (green) and African indicine (pink) ancestries. **B)** Treemix maximum likelihood tree depicting the relationships between taurine cattle breeds (grey: Illumina BovineHD SNP data; black: whole genome data). **C)** Nucleotide diversity in taurine and indicine breeds (Iberian breed names in black).

All analyzed breeds have a positive Tajima’s D (Additional file 1: Figure S4), indicating a reduction in the low-frequency polymorphisms, suggestive of population structure, bias in the choice of genomic markers or of a recent bottleneck probably associated with breeding practices. As observed previously [27], the commercial European breeds have lower nucleotide diversity (average number of pairwise differences) relative to the African breeds (Fig. 2C), potentially a combination of intensive selection and genetic drift resulting from low effective population sizes in European cattle breeds [27]. The Iberian breeds analyzed in this study have, overall, similar or higher values of nucleotide diversity compared to their European counterparts. The lowest values correspond to Mirandesa, Brava and Alentejana, which had been previously shown to have the lowest heterozygosity in a microsatellite panel [19,20]). Management and demographic histories may explain the lower genetic diversity observed in these three breeds. As mentioned above, Mirandesa has recently (since the 1970s) suffered a drastic reduction in population size, and significant inbreeding was detected in Brava and Alentejana [19].

We then used the maximum likelihood approach implemented in Treemix [28] to uncover the historical relationships between the breeds (Fig. 2B). We intersected our whole genome data with the Illumina BovineHD SNP data of 25 European primitive breeds from [4], which shows that our selection of breeds is representative of the Iberian breed context (Fig. 2B). When allowing for one migration event, we observe gene flow from African taurine to the base of the Iberian clade (Fig. 2B) which had been previously suggested to have occurred [14,17,18].

### Iberian cattle show a clear signature of admixture from African cattle and high diversity in mitochondrial DNA and Y chromosome haplotypes

We explicitly test for differential African cattle introgression into Iberian breeds, using the D-statistics [29,30]. We can confirm that there is a significant excess of shared derived alleles in varying amounts between Iberian breeds and the African taurine N’Dama when compared to a panel of European taurine breeds (Fig. 3). This was observed both for southern Iberian Brava that had the largest African (N’Dama) influence, but also in breeds from the north of Portugal such as Barrosã. These results are further corroborated by the occurrence of ~17% of T1-matrilines in the Iberian cattle analyzed here (Fig. 4). The Iberian Peninsula and the Maghreb regions share natural zoo-geographical affinities, and there were complex biogeographic and historic faunal and human relationships during much of the early Holocene, which could explain these patterns of genomic admixture. We did not find evidence of indicine introgression in Iberian cattle, but given that the indicine cattle in our sample has taurine introgression (confirmed by the presence of T1 taurine mitochondrial haplotypes in all the indicine samples of Fig. 4) it is likely that these are not adequate for performing this test.

**Figure 3.**
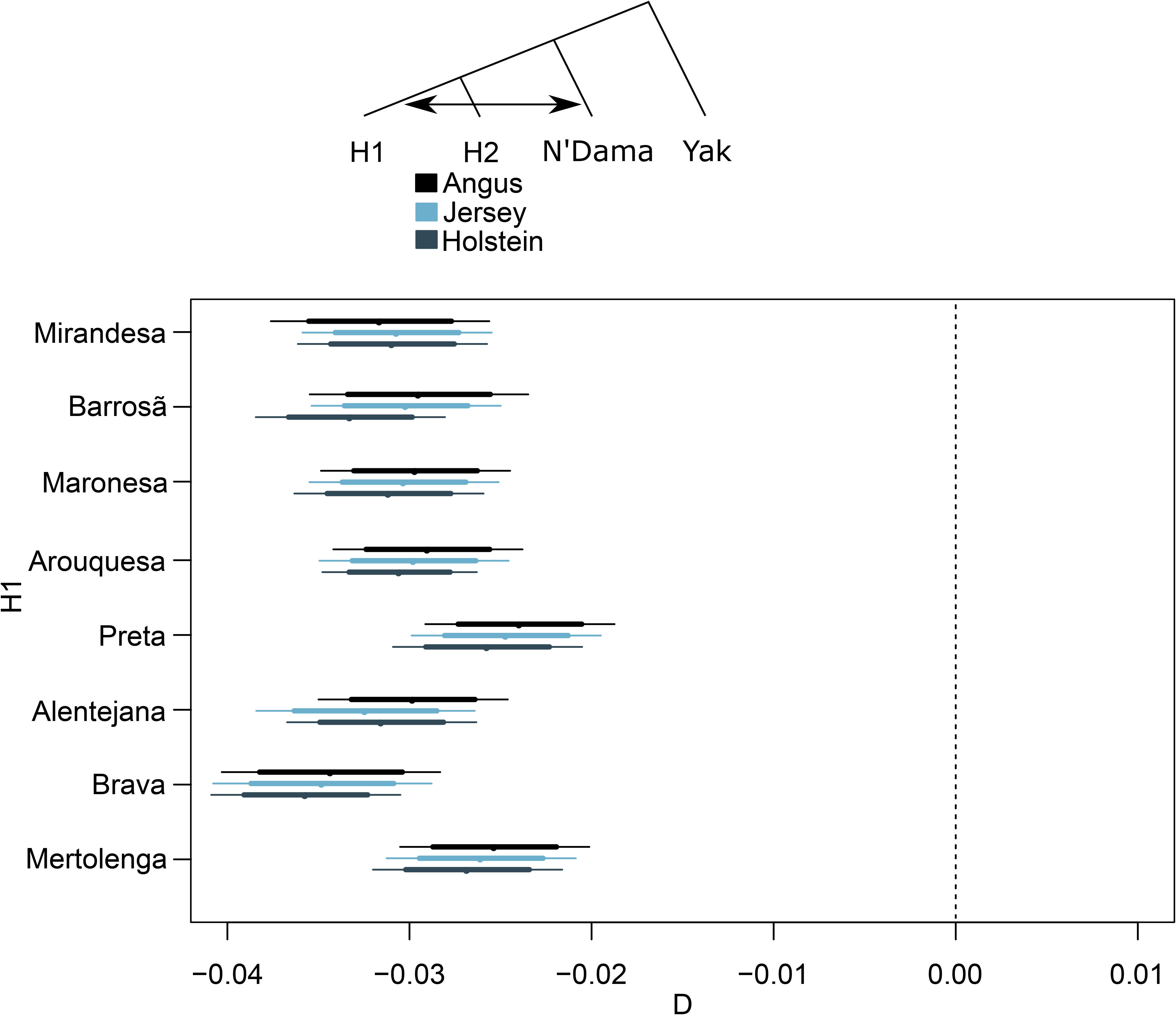
D-statistics determined using genome-wide autosomal data. Negative values indicate an excess of derived alleles shared by the breeds in H1 (denoted in the y-axis) and the African N’Dama breed.

**Figure 4.**
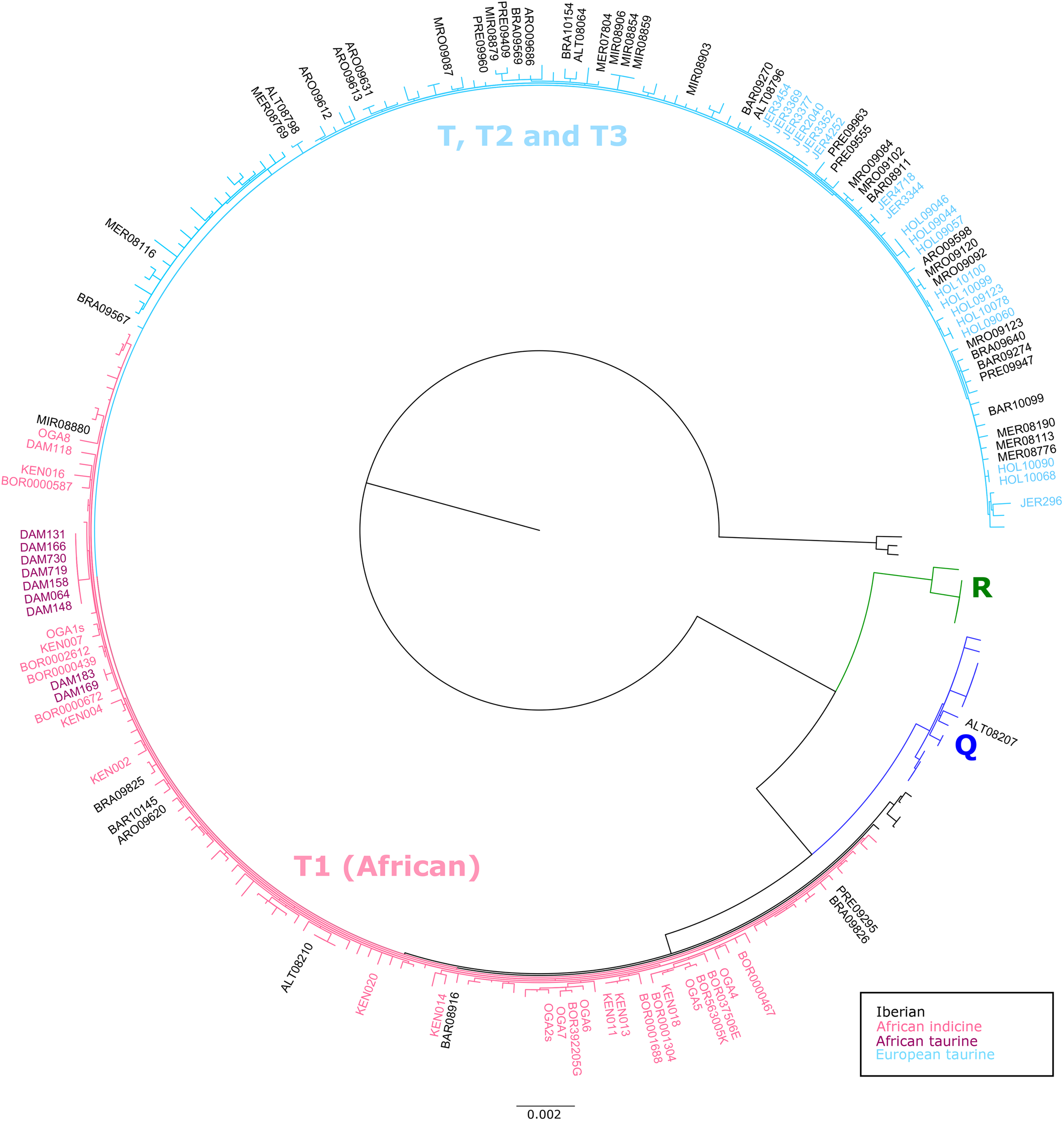
Maximum-likelihood phylogeny of cattle mitogenomes showing that Iberian breeds can be assigned to haplogroups Q, and T, including sub-haplogroup T1 typical of African cattle.

Contrary to previous results [4], we did not find evidence for aurochs introgression into Iberian cattle (Additional file 1: Figure S5) when using sequence data from a 6,750 year-old British aurochs [5]. Given the probable complex population structure of ancient wild cattle in Europe [5,9,31], this result does not preclude that local aurochs introgression occurred, but data from pre-domestic Iberian specimens is required for further testing of this hypothesis.

Because Y-chromosomal variation is geographically structured, with Y1 and Y2 lineages being predominant in northern and central European taurine cattle, respectively, Y-specific markers are useful to investigate crossbreeding [13]. While the Y3 lineage is specific of indicine cattle [14]. In addition, the effective population size of the cattle Y-chromosome is strongly reduced by the reproductive success of popular sires. The paternal diversity of Iberian cattle (Additional file 1: Figure S6 and Table S2) appears to have its origins in the dispersal of a heterogeneous male population since the Neolithic along the Mediterranean route, rather than in the recent admixture of transboundary commercial cattle which are generally fixed for a single patriline (e.g. Holstein-Friesian). Isolation and less intensive selection probably also contributed to preservation of much of the original diversity in this region. Interestingly, Jersey bulls shared a distinct patriline with African cattle (one Ogaden individual; Additional file 1: Figure S6). Previous analyses of Y-chromosome polymorphisms showed that Jersey is fixed for a specific haplotype that is intermediate between Y1 and Y2 haplogroups [14], this may-well represent an African Y-lineage but more comprehensive data from African bulls are needed.

### The impact of breeding practices on chromosomal variation and general patterns of diversification

F_ST_ values between Iberian breeds and other taurine cattle ranged from 12% to 33%, partially overlapping the divergence values observed for comparisons within Iberian breeds (Table 1). Mirandesa, the most divergent within the Iberian breeds, has the highest F_ST_ values relative to all other breeds (Fig. 5A). The taurine breed with the overall highest F_ST_ relative to the Iberian was the Jersey cattle which may be explained by the insular isolated status of this breed [32], although we must note that this might not be a representative sample of the breed.

**Figure 5.**
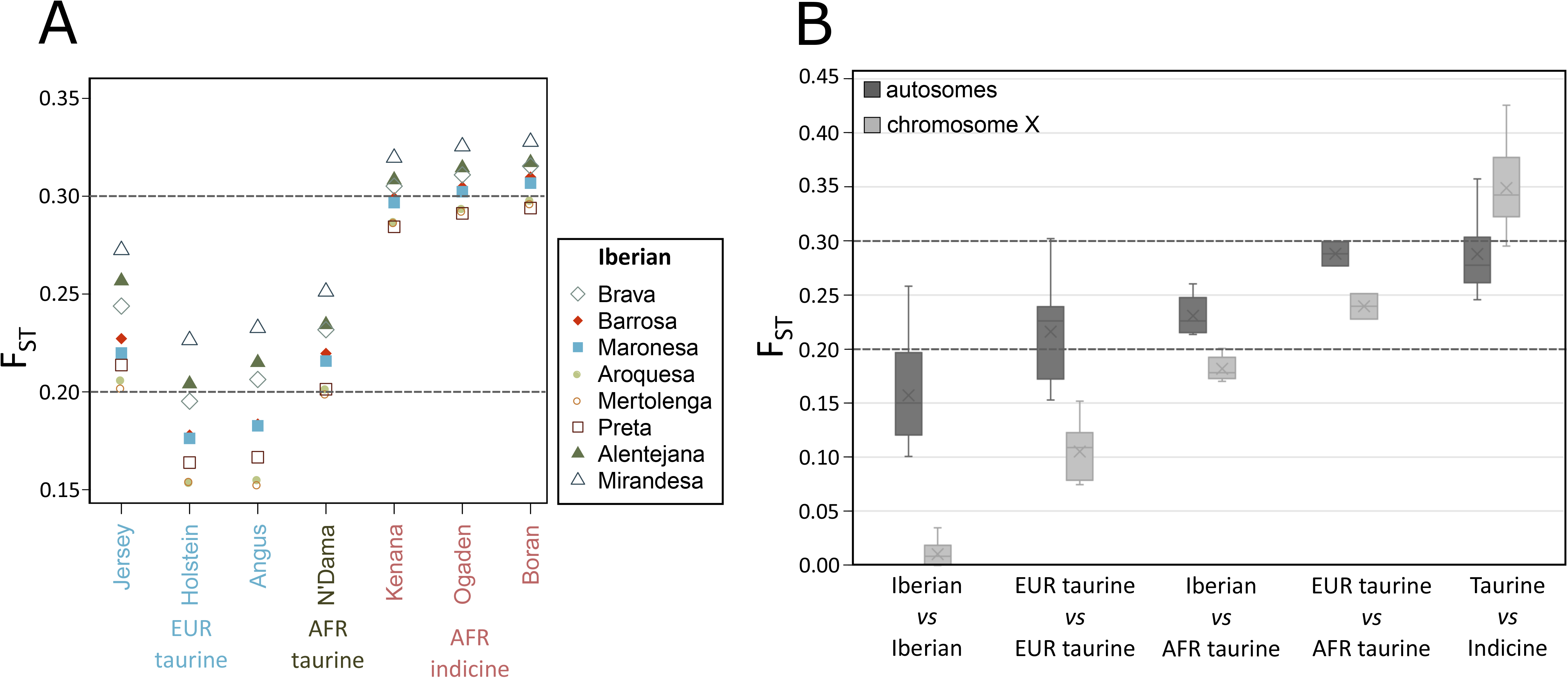
**A)** Autosomal F_ST_ between Iberian cattle and taurine/indicine breeds. **B)** Range of autosomal F_ST_ values for including European taurine (Holstein, Jersey and the Iberian breeds), African taurine (N’Dama), and the African indicine breeds Ogaden, Kenana and Borana. Also shown are the F_ST_ values for sex chromosome X, which is comparatively low within taurine breeds, but shows the expected trend in comparisons with indicine breeds.

The lower effective population size in chromosome X relative to the autosomes should lead to stronger impact of the bottleneck (or population structure) caused by breeding practices, observed in an overall higher Tajima’s (Additional file 1: Figure S4). In this scenario, genetic drift would be expected to result in higher F_ST_ values for chromosome X (lower effective population size [33]) relative to autosomes, which is what we observe when we compare taurine and indicine cattle (Fig. 5B). However, comparisons within taurine and within indicine show a much higher F_ST_ for autosomes than for chromosome X (Fig. 5B; Additional file 1: Figure S7). This is in agreement with extensive male-biased gene flow within taurine and within indicine – since males have a single copy of chromosome X, introgression will be more efficient on the autosomes. It is “known” that female populations are more likely to be geographically constrained and human-driven crossbreeding may have been carried out mainly using males [34]. This could also explain the difference in ancestry assignments for autosomes and chromosome X (Fig. 6), with signatures of previously described indicine admixture in the African taurine autosomes, but not observed in chromosome X.

**Figure 6.**
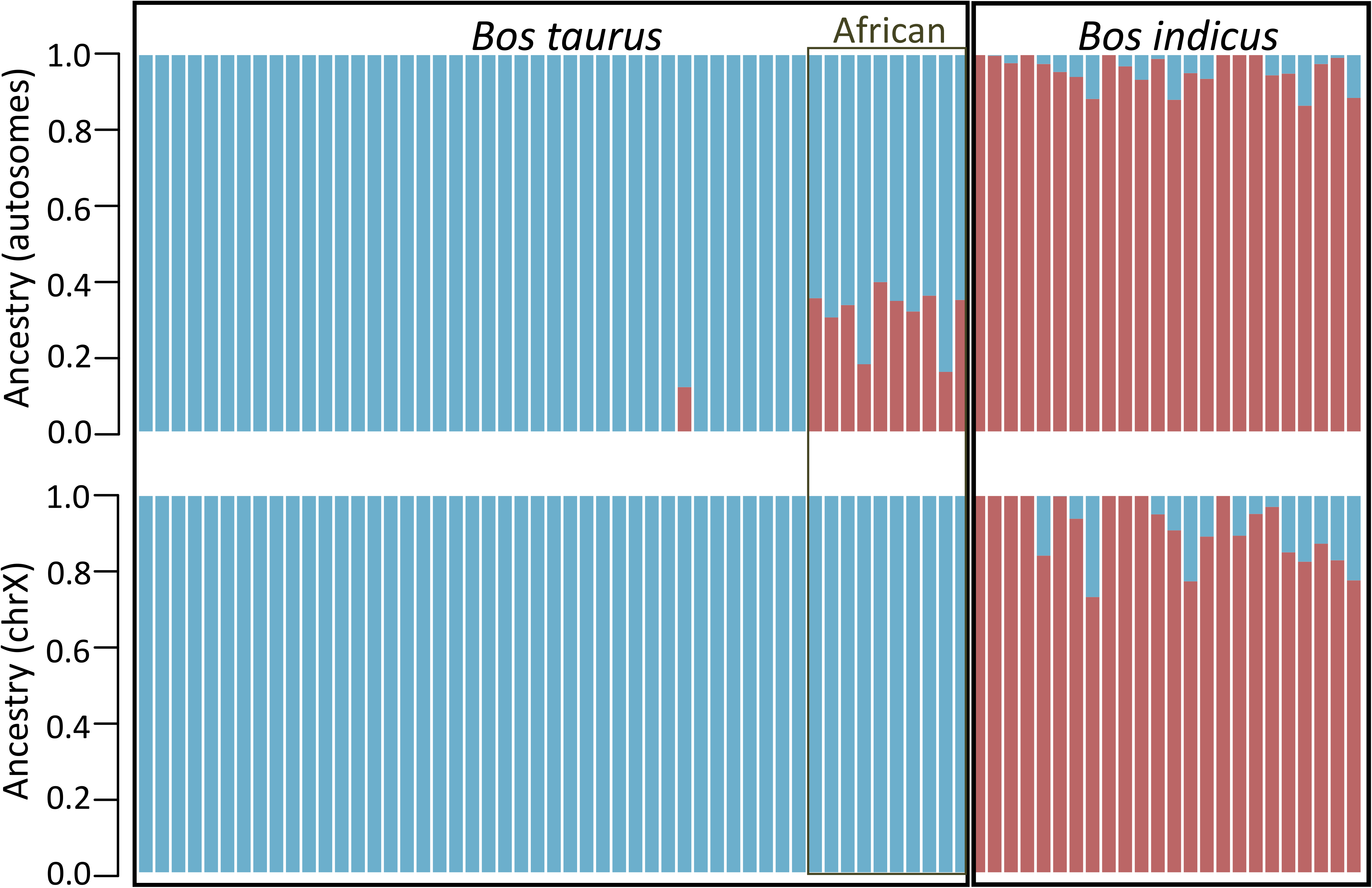
Population structure at K=2 determined using the females individuals only (Table S2). The indicine contribution to African taurine (N’Dama) is not observed in sex chromosome X (bottom) compared to the autosomes (top).

## Conclusion

We release for the first time genomic information on highly diverse peripheral local Iberian cattle, which corroborates these breeds are genetically very distinct in the context of European and African taurine cattle variation. The complex demographic processes underlying the formation of these breeds had profound effects on genomic diversity and resulted in unusual patterns of genetic differentiation for autosomes *vs*. sex chromosomes. Also, Iberian cattle retain much of the original paternal and maternal diversity, which appears to derive from the dispersal of a heterogeneous population since the Neolithic along the Mediterranean route with strong influences from North African taurine cattle, rather than from recent admixture with transboundary commercial cattle. This may have significant impact on the resilience of Iberian cattle to foreseen environmental changes. Not only these breeds produce high-quality certified meat products under local extensive conditions, as they can provide the source for genetic material to improve breeds with depleted genetic diversity, i.e. transboundary commercial cattle. Our results indicate that genetic differentiation measured using chromosome X might be more representative of the native populations of domesticated cattle, and that comparisons between breeds using autosomal data might be misleading without an appropriate demographic model. We also show how whole-genome data are important for uncovering specific patterns related to recent demographic events in breed formation and management, and provide the ground for future studies on the singularity of locally adapted European cattle.

## Materials & Methods

### 1. Materials

Information regarding the breeds and the type of genetic data used to investigate genome diversity and genetic relationships is summarized in supplementary Additional file 1: Table S1. We selected a total of 48 animals representative of Iberian cattle, namely from the Portuguese breeds Alentejana, Arouquesa, Barrosã, Brava de Lide, Maronesa, Mertolenga, Mirandesa and Preta (Fig. 1). The 6 animals of each breed included in our study were nonrelated back to the second generation, originated from several herds, and portray the genetic diversity observed for autosomal microsatellite loci, mitochondrial DNA and Y-chromosome sequences [14,19]. Sampling was done as described in [19], briefly 9 ml of whole-blood were collected from each animal by qualified veterinarians during their routine practice in the framework of official health control programs. Additionally, we used previously generated publicly available genomic data to make population genomics inferences in the context of worldwide cattle: i) shotgun resequencing data of four indigenous African breeds: N’Dama (*Bos taurus*), Ogaden (*Bos indicus*), Boran (*Bos indicus*) and Kenana (*Bos indicus*) [27] (Bioproject ID: PRJNA312138); ii) shotgun resequencing data of three transboundary commercial breeds: Holstein, Jersey, and Angus (Bioproject IDs: PRJNA210521, PRJNA318089 and PRJNA318087, respectively); iii) genotyping Illumina BovineHD SNP data (777,692 SNPs; http://dx.doi.org/10.5061/dryad.f2d1q) of 25 European breeds represented by at least 3 individuals: English Longhorn (England), White Park (England), Galloway (Scotland), Highland (Scotland), Kerry Cattle (Ireland), Heck (Germany), Brown Swiss (Switzerland), Fleckvieh (Switzerland), Dutch Belted (The Netherlands), Dutch Friesian (The Netherlands), Groningen Whiteheaded (The Netherlands), Meuse-Rhine-Yssel (The Netherlands), Busha (Balkan region), Romanian grey (Romania), Boskarin (Check Republic and Hungary), Chianina (Italy), Maremmana (Italy), Maltese (Malta), Cachena (Portugal), Berrenda en Colorado (Spain), Berrenda en negro, (Spain), Cardena (Spain), Lidia (Spain), Limia (Spain), Pajuna (Spain), Sayaguesa (Spain). We also included data of an aurochs (England; Bioproject ID: PRJNA294709) to test for admixture with domesticated cattle. Furthermore, 149 full mitochondrial genomes from NCBI’s PopSets 157778019 [7], 306977267 [35], 355330537 [18], and 946518556 [36] were used together with the mitochondrial consensus sequences obtained from our shotgun data (details below).

### 2. Laboratory procedures

Genomic DNA was extracted using a modified salting-out precipitation method (Gentra Puregene Blood Kit, Qiagen) according to the manufacturer’s recommendations. We prepared equimolar DNA concentrations for all animals before library construction using nanodrop™ and Qubit measurements. Following DNA fragmentation by sonication using a program specific for 550 bp inserts (https://www.diagenode.com/en/p/bioruptor-pico-sonication-device), genomic libraries were prepared using the TruSeq DNA PCR-free Library Preparation Kit (Illumina, San Diego, CA) according to the manufacturer’s protocols. Whole-genome paired-end resequencing data was obtained by pooling 16 samples in each lane and using an Illumina HiSeq1500 instrument with 2×100 bp reads.

### 3. Sequencing data pre-processing

The 48 samples were sequenced to between 1.4X and 2.3X depth of coverage (Additional file 1: Table S3). Methods appropriate for low coverage NGS data [25,37–39] were used throughout the analyses and applied to all samples. Raw Illumina reads were first processed with Trimmomatic [40] for removal of adapter sequences and trimming bases with quality <20 and discarded reads with length <80. Mapping to cattle genome versions UMD_3.1.1 (bosTau8) [23] and Btau_4.6.1 (bosTau7; contains an assembled Y-chromosome) [23], and to the outgroup wild yak (*Bos mutus*; Bioproject ID: PRJNA74739) [24] was done with BWA mem. Reads showing a mapping hit were further filtered for mapping quality >25. PCR duplicates were removed with Picard MarkDuplicates (http://picard.sourceforge.net) and local realignment around indels was done with GATK [41].

### 4. Sequencing error rates

Sequencing error rates were determined in ANGSD [37] using a method that relies on an outgroup and a high quality genome to estimate the expected number of derived alleles (similar to a method described by Reich *et al* [42]). Briefly, if we observe a higher number of derived alleles in an individual we assume that this excess is due to errors. If the high-quality genome is error free, we will obtain an estimate of the true error rate. If there are errors in the high-quality genome, then the estimated error rate can roughly be understood as the excess error rate relative to the error rate of the high-quality genome.

### 5. Population structure

NGSadmix version 32 [25] was used to detect population structure with autosomal data from samples for which shotgun resequencing data was available. NGSadmix infers population structure from genotype likelihoods (that contain all relevant information on the uncertainty of the underlying genotype [43]). NGSadmix was run for K equal to 2, 3, 4, 5 and 6 for sites present in a minimum of 10% of the individuals: a total of 951,213 SNP sites for the 48 Iberian samples (Fig. 1B); 129,829 SNP sites for the data set including all 128 animals (Fig. 2A); 628,774 for SNP sites for the data set including the 94 female individuals (Fig. 6). The program was run with different seed values until convergence was reached.

A principal component analysis using the same SNP set for the Iberian breeds was done with PCAngsd [38] which estimates the covariance matrix for low depth NGS data in an iterative procedure based on genotype likelihoods. Genotype likelihoods for all individuals were generated with ANGSD [37] (options - GL 1 -doGlf 2 -minQ 20 -minMapQ 30).

### 6. Phylogenetic analyses

Treemix [28] was used to infer the admixture graphs (Fig. 2B) using allele counts for 512,358 SNP positions included in the Illumina BovineHD SNP that can be unambiguously assigned to autosomal positions in the cattle reference genome version UMD_3.1.1 [23] using [44]. For shotgun resequencing data, allele counts were obtained from allele frequencies calculated in ANGDS [37] for positions covered in at least 3 individuals. Treemix was run using the global option and standard errors were estimated in blocks with 500 SNPs in each. Even though we do not call genotypes on the shotgun data, the individual breeds where correctly assigned to expected branches in the North/Central European and Iberian clades (Fig. 2B), confirming the robustness of our methodological approach.

The software RAxML [45] version 8.1.7 with 100 rapid bootstrap replicates was used to estimate the phylogenetic trees under the GTR+GAMMA model of sequence evolution for complete mitochondrial sequences from [7,18,35,36] together with consensus sequences from the shotgun resequencing data analyzed in this study obtained by choosing the most common base per position (-doFasta 2 in ANGSD [37]).

### 7. D-statistics

To determine the pattern of excess shared derived alleles between taxa, indicative of introgression, we estimated D-statistics using the wild yak (*Bos mutus*) as an outgroup. All samples were mapped to the yak outgroup genome assembly. The D-statistic [29,30] is approximated by a Gaussian distribution with mean zero [39] in the absence of gene flow between the four populations, allowing for hypothesis testing. We apply an extended version of the D-statistic [39] which can use multiple individuals per population sequenced at low coverage and is implemented in ANGSD [37]. It takes observed allele frequencies for each individual in a population, and then combines them linearly to find an unbiased estimator of population frequency while minimizing the variance [39].

### 8. Assessment of genetic diversity and population differentiation

We used methods based on the site frequency spectrum (SFS) [46,47] to estimate nucleotide diversity, the neutrality test statistic Tajima’s D (Fig. 2C; Additional file 1: Figure S7) and genome-wide F_ST_ values (Fig. 5 and Additional file 1: Figure S7). Briefly, after estimating the SFS, posterior sample allele frequencies are calculated using the global SFS as prior. SFSs estimated separately were used to obtain joint SFSs for population pairs, which are then used to estimate F_ST_. For all pairwise breed comparisons, we determined F_ST_ using autosomes 1 to 29. For comparisons relating to chromosome X, F_ST_ was determined for the sex chromosome and autosomes using only female individuals.

## Declarations

### Ethics approval

Blood samples were collected during routine veterinary checkups in the framework of official health control programs and with the agreement of breeders.

### Consent for publication

Not applicable.

### Data availability

Raw reads are available at https://sid.erda.dk/sharelink/cG0PE8tnjN. The data will be uploaded to the public repository Sequence Read Archive upon acceptance of the manuscript for publication.

### Author contributions

R.R.F. and C.G. designed the study with input from A.E.P. and L.C.; C.G. carried out the sampling and DNA extraction; I.U. and S.A. performed the NGS laboratory work; R.R.F. and C.G. analyzed the data with contributions from E.J.; R.R.F. and C.G. wrote the manuscript with contributions from all authors.

## Acknowledgments

We thank Anders Albrechtsen for stimulating discussion, Nuno Carolino (INIAV, Portugal) for providing information on the demographic characterization of the breeds Alentejana and Brava, and Jolita Dilyté (CIBIO-InBIO) for technical assistance with the Illumina sequencing. The Iberian cattle illustrations were done by biologist Carlos Medeiros for Ruralbit, Portugal (https://ilustracoes.ruralbit.com/). Paulo Gaspar Ferreira (www.paulogasparferreira.com/) and Mariana Bruno-de-Sousa (www.behance.net/MaryAnne_B) were responsible for the graphic concept of the breed distribution figure.

## References

1. Myers N, Mittermeier RA, Mittermeier CG, da Fonseca GAB, Kent J. Biodiversity hotspots for conservation priorities. Nature [Internet]. Nature Publishing Group; 2000 [cited 2017 Jan 4];403:853–8. Available from: http://www.nature.com/articles/35002501

2. Loftus RT, Ertugrul O, Harba AH, El-Barody MA, MacHugh DE, Park SD, et al. A microsatellite survey of cattle from a centre of origin: the Near East. Mol. Ecol. [Internet]. 1999 [cited 2018 Aug 12];8:2015–22. Available from: http://www.ncbi.nlm.nih.gov/pubmed/10632853

3. FAO C on GR for F and AA. The Second Report on the State of the World’s Animal Genetic Resources for Food and Agriculture [Internet]. Scherf BD, Pilling D, editors. Rome; 2015. Available from: www.fao.org/3/a-i4787e.pdf

4. Upadhyay MR, Chen W, Lenstra JA, Goderie CRJ, MacHugh DE, Park SDE, et al. Genetic origin, admixture and population history of aurochs (Bos primigenius) and primitive European cattle. Heredity (Edinb). [Internet]. Nature Publishing Group; 2017 [cited 2018 Apr 26];118:169–76. Available from: http://www.nature.com/articles/hdy201679

5. Park SDE, Magee DA, McGettigan PA, Teasdale MD, Edwards CJ, Lohan AJ, et al. Genome sequencing of the extinct Eurasian wild aurochs, Bos primigenius, illuminates the phylogeography and evolution of cattle. Genome Biol. [Internet]. BioMed Central; 2015 [cited 2018 Apr 30];16:234. Available from: http://genomebiology.com/2015/16/1/234

6. Beja-Pereira A, Caramelli D, Lalueza-Fox C, Vernesi C, Ferrand N, Casoli A, et al. The origin of European cattle: evidence from modern and ancient DNA. Proc. Natl. Acad. Sci. U. S. A. [Internet]. National Academy of Sciences; 2006 [cited 2018 Jul 6];103:8113–8. Available from: http://www.ncbi.nlm.nih.gov/pubmed/16690747

7. Achilli A, Olivieri A, Pellecchia M, Uboldi C, Colli L, Al-Zahery N, et al. Mitochondrial genomes of extinct aurochs survive in domestic cattle. Curr. Biol. [Internet]. Elsevier; 2008 [cited 2018 Apr 18];18:R157–8. Available from: http://www.ncbi.nlm.nih.gov/pubmed/18302915

8. Götherström A, Anderung C, Hellborg L, Elburg R, Smith C, Bradley DG, et al. Cattle domestication in the Near East was followed by hybridization with aurochs bulls in Europe. Proc. Biol. Sci. [Internet]. 2005 [cited 2013 Oct 3];272:2345–50. Available from: http://rspb.royalsocietypublishing.org/content/272/1579/2345.long

9. Mona S, Catalano G, Lari M, Larson G, Boscato P, Casoli A, et al. Population dynamic of the extinct European aurochs: genetic evidence of a north-south differentiation pattern and no evidence of post-glacial expansion. BMC Evol. Biol. [Internet]. BioMed Central; 2010 [cited 2013 Oct 13];10:83. Available from: http://bmcevolbiol.biomedcentral.com/articles/10.1186/1471-2148-10-83

10. Decker JE, McKay SD, Rolf MM, Kim J, Molina Alcalá A, Sonstegard TS, et al. Worldwide patterns of ancestry, divergence, and admixture in domesticated cattle. McVean G, editor. PLoS Genet. [Internet]. Public Library of Science; 2014 [cited 2014 Apr 30];10:e1004254. Available from: http://dx.plos.org/10.1371/journal.pgen.1004254

11. Chen S, Lin B-Z, Baig M, Mitra B, Lopes RJ, Santos AM, et al. Zebu Cattle Are an Exclusive Legacy of the South Asia Neolithic. Mol. Biol. Evol. [Internet]. Oxford University Press; 2010 [cited 2018 Aug 12];27:1–6. Available from: https://academic.oup.com/mbe/article-lookup/doi/10.1093/molbev/msp213

12. Murray C, Huerta-Sanchez E, Casey F, Bradley DG. Cattle demographic history modelled from autosomal sequence variation. Philos. Trans. R. Soc. Lond. B. Biol. Sci. [Internet]. The Royal Society; 2010 [cited 2013 Oct 23];365:2531–9. Available from: http://rstb.royalsocietypublishing.org/content/365/1552/2531.abstract

13. Edwards CJ, Ginja C, Kantanen J, Pérez-Pardal L, Tresset A, Stock F, et al. Dual origins of dairy cattle farming--evidence from a comprehensive survey of European Y-chromosomal variation. Kivisild T, editor. PLoS One [Internet]. Public Library of Science; 2011 [cited 2013 Oct 25];6:e15922. Available from: http://dx.plos.org/10.1371/journal.pone.0015922

14. Ginja C, Penedo MCT, Melucci L, Quiroz J, Martínez López OR, Revidatti MA, et al. Origins and genetic diversity of New World Creole cattle: inferences from mitochondrial and Y chromosome polymorphisms. Anim. Genet. [Internet]. 2010 [cited 2013 Oct 25];41:128–41. Available from: http://www.ncbi.nlm.nih.gov/pubmed/19817725

15. Pelayo R, Penedo MCT, Valera M, Molina A, Millon L, Ginja C, et al. Identification of a new Y chromosome haplogroup in Spanish native cattle. Anim. Genet. [Internet]. 2017 [cited 2018 Jul 6];48:450–4. Available from: http://doi.wiley.com/10.1111/age.12549

16. Beja-Pereira A, Alexandrino P, Bessa I, Carretero Y, Dunner S, Ferrand N, et al. Genetic Characterization of Southwestern European Bovine Breeds: A Historical and Biogeographical Reassessment With a Set of 16 Microsatellites. J. Hered. [Internet]. 2003 [cited 2014 Oct 17];94:243–50. Available from: http://www.ncbi.nlm.nih.gov/pubmed/12816965

17. Colominas L, Edwards CJ, Beja-Pereira A, Vigne J-D, Silva RM, Castanyer P, et al. Detecting the T1 cattle haplogroup in the Iberian Peninsula from Neolithic to medieval times: new clues to continuous cattle migration through time. J. Archaeol. Sci. [Internet]. Academic Press; 2015 [cited 2018 Jul 6];59:110–7. Available from: https://www.sciencedirect.com/science/article/pii/S0305440315001545

18. Bonfiglio S, Ginja C, De Gaetano A, Achilli A, Olivieri A, Colli L, et al. Origin and Spread of Bos taurus: New Clues from Mitochondrial Genomes Belonging to Haplogroup T1. Caramelli D, editor. PLoS One [Internet]. Public Library of Science; 2012 [cited 2018 Jul 6];7:e38601. Available from: http://dx.plos.org/10.1371/journal.pone.0038601

19. Ginja C, Gama LT da, Penedo MCT. Analysis of STR Markers Reveals High Genetic Structure in Portuguese Native Cattle. J. Hered. 2010;101:201–10.

20. Martin-Burriel I, Rodellar C, Cañón J, Cortés O, Dunner S, Landi V, et al. Genetic diversity, structure, and breed relationships in Iberian cattle 1. J. Anim. Sci. 2011;89:893–906.

21. Martin-Burriel I, Rodellar C, Lenstra JA, Sanz A, Cons C, Osta R, et al. Genetic Diversity and Relationships of Endangered Spanish Cattle Breeds. J. Hered. [Internet]. Oxford University Press; 2007 [cited 2018 Jul 6];98:687–91. Available from: https://academic.oup.com/jhered/article-lookup/doi/10.1093/jhered/esm096

22. Albrechtsen A, Nielsen FC, Nielsen R. Ascertainment biases in SNP chips affect measures of population divergence. Mol. Biol. Evol. [Internet]. Oxford University Press; 2010 [cited 2013 Oct 18];27:2534–47. Available from: http://mbe.oxfordjournals.org/content/27/11/2534.long

23. Elsik CG, Tellam RL, Worley KC, Gibbs RA, Muzny DM, Weinstock GM, et al. The genome sequence of taurine cattle: a window to ruminant biology and evolution. Science [Internet]. 2009 [cited 2013 Oct 26];324:522–8. Available from: http://www.sciencemag.org/content/324/5926/522.abstract

24. Qiu Q, Zhang G, Ma T, Qian W, Wang J, Ye Z, et al. The yak genome and adaptation to life at high altitude. Nat. Genet. [Internet]. 2012 [cited 2018 Jul 3];44:946–9. Available from: http://www.ncbi.nlm.nih.gov/pubmed/22751099

25. Skotte L, Korneliussen TS, Albrechtsen A. Estimating individual admixture proportions from next generation sequencing data. Genetics [Internet]. 2013 [cited 2013 Oct 26]; genetics.113.154138-. Available from: http://www.genetics.org/content/early/2013/09/03/genetics.113.154138.abstract?sid=a5f549bf-d0b5-407b-9c13-24142e579370

26. Porter V, Alderson L, Hall SJG, Sponenberg DP. Mason’s World Encyclopedia of Livestock Breeds and Breeding. CABI; 2016.

27. Kim J, Hanotte O, Mwai OA, Dessie T, Bashir S, Diallo B, et al. The genome landscape of indigenous African cattle. Genome Biol. [Internet]. 2017 [cited 2018 Apr 26];18:34. Available from: https://genomebiology.biomedcentral.com/track/pdf/10.1186/s13059-017-1153-y

28. Pickrell JK, Pritchard JK. Inference of population splits and mixtures from genome-wide allele frequency data. Tang H, editor. PLoS Genet. [Internet]. Public Library of Science; 2012 [cited 2013 Oct 24];8:e1002967. Available from: http://dx.plos.org/10.1371/journal.pgen.1002967

29. Patterson NJ, Moorjani P, Luo Y, Mallick S, Rohland N, Zhan Y, et al. Ancient admixture in human history. Genetics [Internet]. 2012; Available from: http://www.genetics.org/content/early/2012/09/06/genetics.112.145037.abstract

30. Reich D, Thangaraj K, Patterson N, Price AL, Singh L. Reconstructing Indian population history. Nature [Internet]. 2009 [cited 2018 Apr 17];461:489–94. Available from: http://www.ncbi.nlm.nih.gov/pubmed/19779445

31. Scheu A, Powell A, Bollongino R, Vigne J-D, Tresset A, Çakırlar C, et al. The genetic prehistory of domesticated cattle from their origin to the spread across Europe. BMC Genet. [Internet]. BioMed Central; 2015 [cited 2018 Aug 12];16:54. Available from: http://www.biomedcentral.com/1471-2156/16/54

32. Chikhi L, Goossens B, Treanor A, Bruford MW. Population genetic structure of and inbreeding in an insular cattle breed, the Jersey and its implications for genetic resource management. Heredity (Edinb). [Internet]. 2004 [cited 2018 Aug 12];92:396–401. Available from: http://www.ncbi.nlm.nih.gov/pubmed/15014423

33. Wilson Sayres MA. Genetic Diversity on the Sex Chromosomes. Genome Biol. Evol. [Internet]. Oxford University Press; 2018 [cited 2018 Apr 19];10:1064–78. Available from: https://academic.oup.com/gbe/article/10/4/1064/4895090

34. Lenstra J, Ajmone-Marsan P, Beja-Pereira A, Bollongino R, Bradley D, Colli L, et al. Meta-Analysis of Mitochondrial DNA Reveals Several Population Bottlenecks during Worldwide Migrations of Cattle. Diversity [Internet]. Multidisciplinary Digital Publishing Institute; 2014 [cited 2018 Aug 12];6:178–87. Available from: http://www.mdpi.com/1424-2818/6/1/178

35. Bonfiglio S, Achilli A, Olivieri A, Negrini R, Colli L, Liotta L, et al. The Enigmatic Origin of Bovine mtDNA Haplogroup R: Sporadic Interbreeding or an Independent Event of Bos primigenius Domestication in Italy? Kivisild T, editor. PLoS One [Internet]. Public Library of Science; 2010 [cited 2018 Apr 18];5:e15760. Available from: http://dx.plos.org/10.1371/journal.pone.0015760

36. Olivieri A, Gandini F, Achilli A, Fichera A, Rizzi E, Bonfiglio S, et al. Mitogenomes from Egyptian Cattle Breeds: New Clues on the Origin of Haplogroup Q and the Early Spread of Bos taurus from the Near East. Caramelli D, editor. PLoS One [Internet]. 2015 [cited 2018 Aug 22];10:e0141170. Available from: http://www.ncbi.nlm.nih.gov/pubmed/26513361

37. Korneliussen TS, Albrechtsen A, Nielsen R. ANGSD: Analysis of Next Generation Sequencing Data. BMC Bioinformatics [Internet]. BioMed Central Ltd; 2014 [cited 2015 Jul 31];15:356. Available from: http://www.biomedcentral.com/1471-2105/15/356

38. Meisner J, Albrechtsen A. Inferring Population Structure and Admixture Proportions in Low Depth NGS Data. bioRxiv [Internet]. Cold Spring Harbor Laboratory; 2018 [cited 2018 Jun 16];302463. Available from: https://www.biorxiv.org/content/early/2018/05/23/302463.figures-only

39. Soraggi S, Wiuf C, Albrechtsen A. Powerful Inference with the D-Statistic on Low-Coverage Whole-Genome Data. G3 (Bethesda). [Internet]. G3: Genes, Genomes, Genetics; 2018 [cited 2018 Apr 17];8:551–66. Available from: http://www.ncbi.nlm.nih.gov/pubmed/29196497

40. Bolger AM, Lohse M, Usadel B. Trimmomatic: a flexible trimmer for Illumina sequence data. Bioinformatics [Internet]. Oxford University Press; 2014 [cited 2016 Aug 11];30:2114–20. Available from: http://www.ncbi.nlm.nih.gov/pubmed/24695404

41. DePristo MA, Banks E, Poplin R, Garimella K V, Maguire JR, Hartl C, et al. A framework for variation discovery and genotyping using next-generation DNA sequencing data. Nat. Genet. [Internet]. Nature Publishing Group, a division of Macmillan Publishers Limited. All Rights Reserved.; 2011 [cited 2014 Jan 21];43:491–8. Available from: http://dx.doi.org/10.1038/ng.806

42. Reich D, Green RE, Kircher M, Krause J, Patterson N, Durand EY, et al. Genetic history of an archaic hominin group from Denisova Cave in Siberia. Nature [Internet]. Nature Publishing Group, a division of Macmillan Publishers Limited. All Rights Reserved.; 2010 [cited 2014 Jan 22];468:1053–60. Available from: http://dx.doi.org/10.1038/nature09710

43. da Fonseca RR, Albrechtsen A, Themudo GE, Ramos-Madrigal J, Sibbesen JA, Maretty L, et al. Next-generation biology: Sequencing and data analysis approaches for non-model organisms. Mar. Genomics [Internet]. 2016 [cited 2016 May 14];30:3–13. Available from: http://www.sciencedirect.com/science/article/pii/S1874778716300368

44. Nicolazzi EL, Caprera A, Nazzicari N, Cozzi P, Strozzi F, Lawley C, et al. SNPchiMp v.3: integrating and standardizing single nucleotide polymorphism data for livestock species. BMC Genomics [Internet]. 2015 [cited 2018 Jun 29];16:283. Available from: http://www.ncbi.nlm.nih.gov/pubmed/25881165

45. Stamatakis A. RAxML version 8: a tool for phylogenetic analysis and post-analysis of large phylogenies. Bioinformatics [Internet]. 2014;30:1312–3. Available from: http://www.pubmedcentral.nih.gov/articlerender.fcgi?artid=3998144&tool=pmcentrez&rendertype=abstract

46. Korneliussen TS, Moltke I, Albrechtsen A, Nielsen R. Calculation of Tajima’s D and other neutrality test statistics from low depth next-generation sequencing data. BMC Bioinformatics [Internet]. 2013 [cited 2013 Oct 21];14:289. Available from: http://www.biomedcentral.com/1471-2105/14/289

47. Nielsen R, Korneliussen T, Albrechtsen A, Li Y, Wang J. SNP calling, genotype calling, and sample allele frequency estimation from new-generation sequencing data. PLoS One [Internet]. Public Library of Science; 2012;7:e37558. Available from: http://dx.doi.org/10.1371%252Fjournal.pone.0037558

